# Characterization of CRISPR/Cas9 RANKL knockout mesenchymal stem cell clones based on single-cell printing technology and emulsion coupling assay as a low-cellularity workflow for single-cell cloning

**DOI:** 10.1101/2020.08.17.253559

**Authors:** Tobias Groß, Csaba Jeney, Darius Halm, Günter Finkenzeller, G. Björn Stark, Roland Zengerle, Peter Kolaty, Stefan Zimmermann

## Abstract

The homogeneity of the genetically modified single-cells is a necessity for many applications such as cell line development, gene therapy, and tissue engineering and in particular for regenerative medical applications. The lack of tools to effectively isolate and characterize CRISPR/Cas9 engineered cells is considered as a significant bottleneck in these applications. Especially the incompatibility of protein detection technologies to confirm protein expression changes without a preconditional large-scale clonal expansion, creates a gridlock in many applications. To ameliorate the characterization of engineered cells, we propose an improved workflow, including single-cell printing/isolation technology based on fluorescent properties with high yield, a genomic edit screen (surveyor assay), mRNA rtPCR assessing altered gene expression and a versatile protein detection tool called emulsion-coupling to deliver a high-content, unified single-cell workflow. The workflow was exemplified by engineering and functionally validating RANKL knockout immortalized mesenchymal stem cells showing altered bone formation capacity of these cells. The resulting workflow is economical, without the requirement of large-scale clonal expansions of the cells with overall cloning efficiency above 30% of CRISPR/Cas9 edited cells. Nevertheless, as the single-cell clones are comprehensively characterized at an early, highly parallel phase of the development of cells including DNA, RNA, and protein levels, the workflow delivers a higher number of successfully edited cells for further characterization, lowering the chance of late failures in the development process.

**Author summary:** I completed my undergraduate degree in biochemistry at the University of Ulm and finished my master's degree in pharmaceutical biotechnology at the University of Ulm and University of applied science of Biberach with a focus on biotechnology, toxicology and molecular biology. For my master thesis, I went to the University of Freiburg to the department of microsystems engineering, where I developed a novel workflow for cell line development. I stayed at the institute for my doctorate, but changed my scientific focus to the development of the emulsion coupling technology, which is a powerful tool for the quantitative and highly parallel measurement of protein and protein interactions. I am generally interested in being involved in the development of innovative molecular biological methods that can be used to gain new insights about biological issues. I am particularly curious to unravel the complex and often poorly understood protein interaction pathways that are the cornerstone of understanding cellular functionality and are a fundamental necessity to describe life mechanistically.

## Introduction

There is a high demand for well-characterized genetically engineered single-cell clones [1]. The assurance of their clonality and lineage traceability are important not only for pharmaceutical but also for cell therapeutic applications particular for regenerative medical uses [2], which is also enforced by the regulatory requirements of the European Medicine Agency (EMA) and the Food and Drug Administration (FDA) [3, 4]. This work aims to provide an improved, more parallel workflow, without time-consuming clonal expansion, to generate deeply characterized single-cell clones that can meet these quality requirements.

Most genetic engineering methods such as CRISPR/Cas9 are error-prone, generating a non-homogeneous population of cells by failing to introduce the engineered changes correctly, having off-targets, monoallelic modifications, and many non-edited cells [5], rendering the clonal isolation of the cells and the characterization of the clones mandatory before their use. Nevertheless, as the genomically correct modification does not ensure the intended gene expression changes, the validation of the requirements necessitates a high-content characterization at genomic, mRNA expression, and also protein level. To fulfill these analytical needs a battery of technologies is applied, which introduce their own, in many cases, disparate requirements. As a consequence, the current approaches frequently include the costly, failure-prone, and time-consuming expansion of the cells solely to provide material for the subsequent analytical methods. This is especially true for protein analytic technologies, as they are in regard to most frequently used methods, and in contrast to genomic and RNA expression technologies, not molecularly sensitive. While genomic and RNA expression detection technologies can use even a single cell of sample for their analysis [6], protein analytics need several magnitude larger sample amounts. The available classic methods such as mass flow cytometry, mass spectrometry, ELISA, or Western Blot often require a large number of cell material in order to detect the targeted protein [7–10]. However, new protein analytical technologies are emerging such as proximity ligation assay [11], proximity extension assay [12], or single-cell mass spectrometry [13].

Recently, single-cell printing technology (SCP) is emerging from others [14, 15] as a gentle, low-cost and highly controlled technology, applicable for a wide array of specific cell-cloning applications ranged from the 0.8 *μ*m prokaryotes to 100 *μ*m plant cells [16, 17]. It is based on inject-jet-like technologies generating on-demand free-flying microdroplets encapsulating cells to deposit them on variable substrates using a non-contact dispensing process. The technology is described in more detail by *Gross et al*. 2013.

SCP can be considered as a very gentle single-cell isolation technology that is comparable to manual pipetting in terms of the applied shear stress [18]. SCP is also superior to fluorescence-associated cell sorting (FACS), which is also frequently used for single-cell isolation, as FACS has a negative effect on cloning efficiency of sensitive or partly-damaged cells (e.g. by transfection) by applying high shear forces and electrostatic charging of cells [15]. SCP is especially well-suited for cloning of engineered cells, besides preserving their viability, it provides direct proof of single-cell clonality (e.g. delivers trustable records of each printed cell), has both native and fluorescent sorting capabilities, and can be coupled to single-cell analytical pipelines. Recently, SCP has become a frequent choice for single-cell isolation purposes in different workflows including subsequent analytics [19].

As discussed above, genomic and transcriptomic assays usually do not pose significant issues but the mandatory protein analytics are a central challenge in cloning workflows. Albeit the recent developments of sensitive protein assays and their wide variety of applications further improvement is still necessary. An ideal assay has a homogenous measurement principle (e.g. no washing steps), can be used for sample sizes with very low cell numbers down to a few dozen cells, and yet provides highly sensitive and quantitative detection of many proteins parallel, including protein post-translational modifications, and protein-protein interactions. Recently, emulsion coupling [20] promises to fulfill these requirements.

Emulsion coupling is based on a digital method that is similar to the droplet digital PCR (ddPCR) [21, 22] and detects the partitioning differences of two DNA oligonucleotide labeled antibodies per target in the presence of target protein compared to their free, unbound distribution in ddPCR emulsion. This assay is homogenous, molecularly sensitive, quantitative, and due to its two antibody-principle, it is highly specific and can be read by sequencing (e.g. by next-generation sequencing) in a highly parallel way. Its low sample requirement and parallelity make it an ideal candidate to establish a cell cloning workflow with minimum clonal expansion.

To provide an experimental proof of concept, immortalized mesenchymal stem cells have been genetically engineered to improve their bone-forming capacity for potential regenerative medical applications. For this, Tumor Necrosis Factor Superfamily Member 11 (*TNFSF11*) was knocked out as suggested among others by *Walsh and Choi 2014* [23–25], which encodes the receptor activator for nuclear factor kappa B ligand (RANKL) [26]. RANKL has been shown to play a crucial role in bone homeostasis by orchestrating the balance between bone-generating osteoblasts and bone-degrading osteoclasts [27–29] via the so-called OPG/RANKL/RANK pathway [30]. Briefly, RANKL is expressed in bone tissue by mesenchymal stem cells (MSCs), osteoblasts, and T-cells, among others [31]. In the presence of RANKL, the receptor activator for nuclear factor kappa B (RANK) is activated which stimulates pre-osteoclasts to differentiate into osteoclasts which in turn degrade bone [32, 33]. For bone formation, MSCs differentiate into osteoblasts and deposit calcified structures [34]. By *TNFSF11* knockout, the genetically modified MSCs and their progenitors can no longer recruit osteoclasts and as a consequence, the bone formation at the side of their implantation in a regenerative medical application could be potentially improved.

An improved workflow was provided including SCP technology with fluorescent sorting, and limited clone expansion to enable genomic edit screen (surveyor assay), mRNA RT-PCR assessing altered gene expression, and emulsion coupling to deliver a high-content, unified single-cell cloning workflow.

## Materials and methods

### Cell culture

Immortalized MSCs were kindly provided by Prof. Dr. Matthias Schieker from the Laboratory of Experimental Surgery and Regenerative Medicine of the Ludwig-Maximilians-University of Munich. They were immortalized by retroviral transduction of human telomerase reverse transcriptase (hTERT MSC) [35]. The cells were cultivated in Nunc™ EasYFlask™ Nunclon™ Delta Surface (Thermo Scientific) using the cell culture medium MEM alpha Medium (1x) + GlutaMax™ - I (Gibco), 10% FBS (Gibco) and 1% Pen-Strep (Gibco) at 37°C/5% CO_2_.

### Determination of the cumulative population doublings

The population doubling (PD) was determined by tracking the cell number of the initially seeded cells (y) and the cell yield during cell passaging (x). The PD was calculated by the following formula: PD=(ln(x) - ln(y))/ln(2). The cumulative population doubling (CPD) is the total number of times the cells in a given population have doubled in culture, which was plotted against the days after single-cell isolation.

### Plasmids

The RANKL specific gRNA construct (pNV-RANKL/KO) based on the parenteral gRNA-Cas9-2A-GFP vector, which is transiently expressing gRNA and Cas9 and enables GFP selection of transfected MSCs, was purchased from abm (Richmond, Canada). Three vectors were purchased with the following gRNA sequences: pNV-RANKL1/KO - 5’-CAGGAATTACAACATATCGT-3’ (472221110290), pNV-RANKL2/KO - 5’-CAGCGATGGTGGATGGCTCA-3’ (472221110390), pNV-RANKL3/KO - 5’-TTAATAGTGAGATGAGCAAA-3’ (472221110490). pmaxGFP (purchased from Lonza, Human MSC Nucleofector^®^ Kit) was used to control transfection efficiency. Plasmids were propagated by transforming *Escherichia Coli* (Mix&Go! Competent Cells - DH5 Alpha, Zymo Research) using standard procedures. The plasmids were purified using the ZymoPURE™ II Plasmid Midiprep Kit (Zymo Research) according to the manufacturer’s instructions. The plasmid concentration was measured with the NanoDrop™ One. The plasmids were stored at – 20°C.

### Transfection

Sub-confluent hTERT MSCs were trypsinized, harvested, and washed by PBS using standard procedures. For nucleofection of hTERT MSC cells, cells were resuspended in 100 *μ*l Human MSC Nucleofector Solution (Human MSC Nucleofector^®^ Kit - Lonza) and the transfection was carried out according to the manufacturer’s instructions using different amount of gRNA plasmids or pmaxGFP plasmid. The transfected cells were incubated at 37°C/5% CO_2_ for 24 h before further steps. Transfection efficiencies were determined using the data obtained during single-cell printing. GFP expressing cells were set relative to the total count of detected cells.

### Single-cell printing and single-cell cloning

The cell samples were prepared for single-cell printing according to *Riba et al. 2018*. The f.sight^®^ single-cell printer - SCP (cytena) selected and printed the GFP-expressing cells according to their fluorescent signal and sorted them according to the morphological criteria of - size (15-30 *μ*m) and roundness (0.6 - 1). The cells were printed into wells of a 96 well-plate (flat bottom clear polystyrene, Greiner) containing 200 *μ*l cell culture medium (see cell culture). After printing, the printed cells were detected with the high-resolution imager NyOne™ (SynenTec) either in single-cell colony (SCC) or in fluorescence-activated SCC (FASCC) mode. The single-cell printing efficiency was determined based on *Gross et al. 2013*. Briefly, images that were taken of the printed cells during the printing process were manually analyzed for single, multiple, void, or uncertain printing events. The percentage of each event was calculated relative to the total printing events. The single-cells cloning efficiency was accessed by the high-resolution imager NyOne™ (SynenTec) using its confluence mode. Cloning efficiency was determined by calculating the percentage of the wells with cell colonies relative to the total count of printed wells with single-cells.

### Genome editing detection

Genomic DNA was extracted from 8.0×10^5^ hTERT MSCs with the Quick-DNA™ Microprep Plus Kit (Zymo Research) and then analyzed for indel mutations by the surveyor assay using the Alt-R™. Genome Editing Detection kit (IDT) according to the manufacturer’s instructions. Briefly, the genomic DNA was extracted from the samples and the loci of the edited sites were amplified by PCR of both. Amplifications were carried out using the HotStarTaq Plus Master Mix Kit (Qiagen) according to the manufacturer’s instructions. Unmodified hTERT MSC reference with CRISPR edited DNA was combined to form heteroduplexes in a 1:1 ratio, 5 *μ*l unmodified hTERT MSC reference with 5 *μ*l CRISPR edited DNA. The heteroduplexes were digested according to the Alt-R™ Genome Editing Detection Kit. Controls A and B from the Alt-R™ Genome Editing Detection Kit were used and treated in the same way. 4 *μ*l of the digested DNA was mixed with 2 *μ*l 1:100 in DMSO diluted GelRed™ (Biotium) dye and loaded onto a 1% agarose gel and as a ladder the 100 bp plus DNA ladder from VWR was used. Gel images were taken by the camera system INGenius (Syngene). The forward primer 5’-AAGTTCTGCGGCCCAGTTTA-3’ and reverse primer 5’-AGGGAGAGAAAGGAACCTCTG-3’ were used. PCR conditions having a hot-start at 95°C for 5 min, with cycling conditions at 95°C for 30 sec, 30 sec annealing temperature at 64’C, and with an extension time of 1.5 min at 72’C and 40 cycle rounds with a final extension time at 72’C for 10 min. Control A/B shows three bands at 690 bp, 463 bp, and 256 bp for digested heteroduplexes, control A for a homoduplex at 690 bp, genomic DNA from unmodified hTERT MSCs at 1850 bp and the negative control (-) no signal.

### Real-time quantitative PCR

The gene expression level of *TNFSF11* in hTERT MSCs and the RANKL-KO hTERT MSCs were determined using qPCR. Briefly, total RNA was isolated from 5 × 10^5^ cells with the AllPrep© DNA/RNA/Protein Mini Kit (Qiagen) according to the manufacturer’s instructions. Two assays were used simultaneously with a VIC-probe located at the 5’ end of the *TNFSF11* transcript (Hs00243519_m1 TaqMan assay) and a FAM-probe located at the 3’ end of the *TNFSF11* transcript (Hs00243522_m1 TaqMan assay). The qPCR was performed using the TaqMan™ RNA-to-CT™ 1-Step kit (Thermo Fischer Scientific) and was run in a RotorGene-6000-2-plex (Qiagen). PCR conditions having reverse transcription at 48’C for 15 min, hot-start at 95°C for 10 min, with cycling conditions starting with a denaturation step at 95’C for 15 sec, anneal/extend step at 60’C for 1 min. The results were analyzed with the Rotor-Gene Q Series Software from Qiagen.

### Emulsion Coupling Assay

10^6^ hTERT MSCs, and different RANKL-KO (see section Plasmids) hTERT MSCs were harvested and lysed and processed according to the emulsion coupling protocol described by *Karakus et al. 2019* and is briefly described below. Two anti-RANKL antibodies, RANKL-G1 (Santa Cruz Biotechnology) FAM signal, anti-CD254 (Biolengend) VIC signal, were used. The oligonucleotide labeling of the antibodies was carried out according to *Karakus et al. 2019*. Lysed cells were incubated with the antibodies overnight and diluted 100,000-fold. Emulsification, ddPCR, and readout were performed using the QX200 Droplet Digital™ PCR System (Bio-Rad) following standard ddPCR protocol. As a negative control, the antibodies were mixed without antigen and processed in the same way as the samples (antibody-binding-control, ABC). The amount of protein RANKL was measured by detecting the partitioning differences induced by concurrent binding of the antibodies to RANKL. The measurements were normalized against the ABC control, which defines the zero level detection by the ABC signal.

### Osteogenic differentiation

10^4^ cells of the CRISPR-edited RANKL knockout cell clones and unmodified hTERT MSCs were seeded in 6-well plates. They were incubated in 2 ml of osteogenic differentiation medium (DMEM low glucose (Gibco), 10% FBS, 1%PenStrep, 100 nM dexamethasone, 50 *μ*g/ml ascorbate, 10 mM ß-glycerophosphate) for 7, 14 and 21 days. In order to stain calcified deposits, 2 g of Alizarin Red (Sigma-Aldrich) was dissolved in 100 ml distilled water and the pH was adjusted to 4.5 – 5 by NaOH and the solution was filtered. Before staining, the culture medium was removed from the cells and they were washed 2 times with PBS. The cells were then fixed with 4% formalin in PBS for 10 minutes and washed 2 times with PBS. For staining, 2 ml Alizarin Red staining solution was added to the wells, and the cells were stained in the dark for 45 minutes. The staining solution was removed and the cells were carefully washed 5 times with PBS. Images of the stained cells were captured with the microscope DMi1 (Leica).

## Results and Discussion

### Experimental design

Working on the generation of genetically engineered cells raises a number of problems, among others the generation of well-characterized clones and their assurance and documentation of clonality. We propose a practicable workflow for single-cell cloning addressing these problems which additionally provides low-cellularity analytics. The individual steps of the improved workflow are displayed in Fig1. To characterize a large number of clones in economical timescales without excessive clonal expansion, PCR based technologies are available. However, such technologies are highly limited in protein analytics. We devised a technological solution to overcome this limitation: (i) modifications at the genomic level were detected using a surveyor assay, (ii) transcription levels of target genes were checked by RT-PCR and (iii) target protein expression by the clones was measured by emulsion coupling. All these assays enable the characterization of the clones from a few hundred to thousand cells as sample amount (low-demand assay - LDA), fostering low clonal expansion and high throughput characterization, but still providing high-content information about the clones, facilitating successful functional assays of the cherry-picked clones. This workflow is compatible with industrial requirements, highly automatable, and can easily be transferred to other cell types and genetic targets. For a proof of concept, the workflow was applied on hTERT MSCs which were *TNFSF11* edited using the CRISPR/Cas9 system [36], to demonstrate the efficient single-cell handling and sorting of genetically modified cells using SCP as well as accessing the performance of the LDAs in this environment.

**Fig 1.**
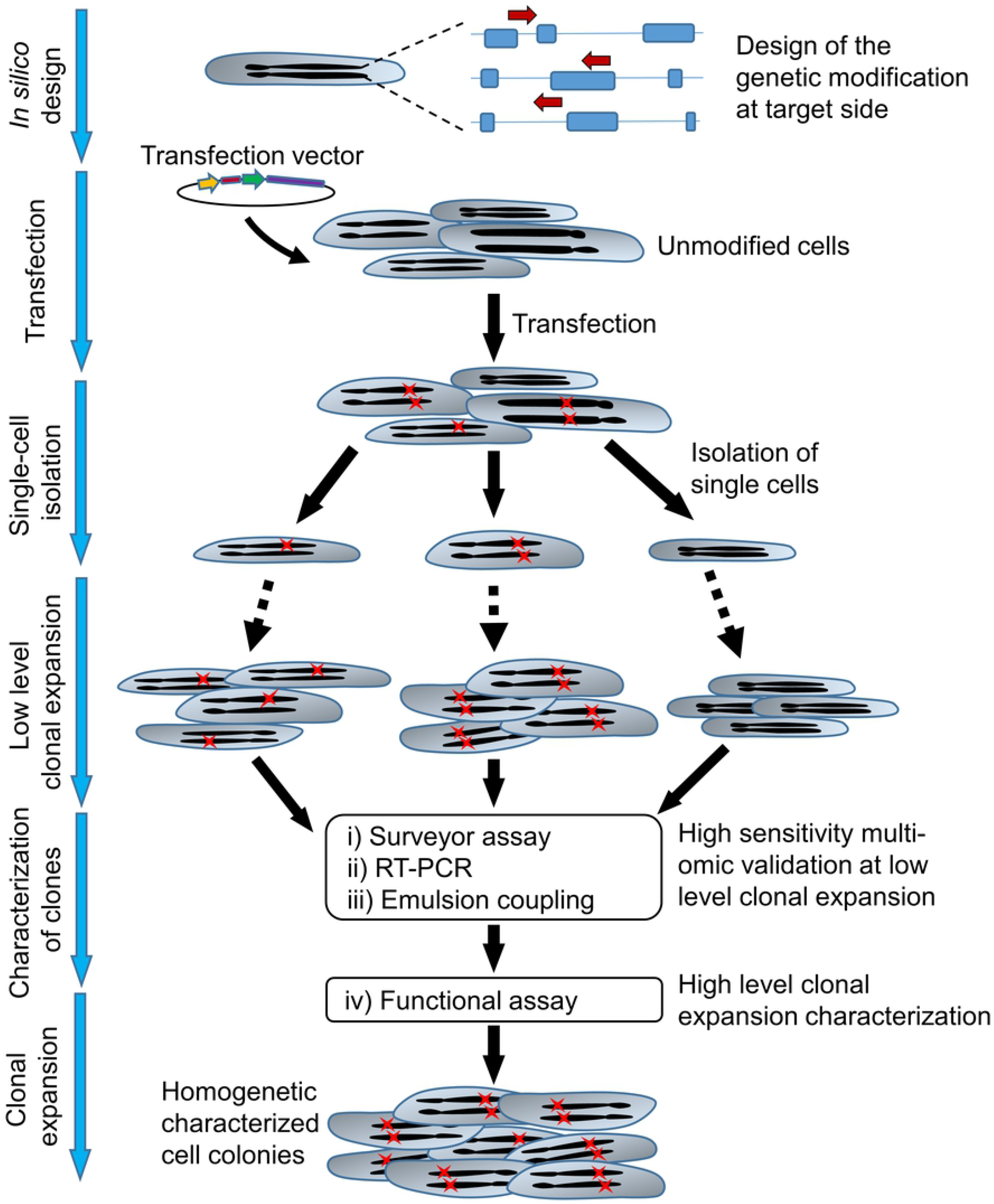
Improved workflow to establish and characterize CRISPR/Cas9 edited cells with minimal clonal expansion. gRNAs transfected cells were isolated and sorted by SCP detecting GFP signal. To characterize the introduced genetic changes, RNA, and protein expressions, a set of assays requiring a low number of cells was proposed: (i) surveyor assay (DNA edits), (ii) RT-PCR (RNA expression) and (iii) emulsion coupling (protein expression). The selected clones are ready for further confirmatory assays (iv). Arrows indicate gRNAs, red crosses symbolize introduced genetic changes.

### CRISPR/Cas9 gene editing and single-cell cloning

hTERT MSCs were transfected with the pNV-RANKL/KO plasmids, CRISPR/Cas9 knockout vectors, with a GFP marker gene containing different gRNAs directed against *TNFSF11* (see Material and Methods). In order to sort transfected single-cells from the untransfected ones, the SCP technology with fluorescent sorting (cytena, f.sight™) was applied by detecting and sorting cells by the GFP signal of the transfected constructs. In Fig 2A, exemplary five images of the isolation process of a transfected and GFP expressing hTERT MSC are shown. As a control, hTERT MSCs were transfected with pmaxGFP to evaluate the efficiency of printing, sorting, and cloning and to optimize the process. hTERT MSCs were transfected with pNV-RANKL/KO gRNA variants and printed and sorted according to the determined parameters with regard to size, roundness and laser intensity, exposure time, and fluorescent threshold. Although the hTERT MSCs were transfected with different transfection efficiencies, pmaxGFP had an efficiency of 61% and pNV-RANK/KO only 6%, and additionally, pNV-RANK/KO transfection induced significantly more dead cells, at the same time a cloning efficiency of 30% was achieved with the transfected cells. The differences in transfection efficiency can be explained by the smaller size of the pmaxGFP control vector, which is approximately 3.5 kb compared to 12 kb of the pNV-RANKL/KO plasmid [37]. The pmaxGFP plasmid was designed to have high-intensity GFP expression while the SFFV promoter in the pNV-RANKL/KO vector transcribed a Cas9-GFP multicistronic transcription unit, resulting in a weaker signal. We also noted a minimal difference in the cloning efficiency of transfected hTERT MSCs (31.3 ± 8.0%) and non-transfected hTERT MSCs (39.6 ± 15.6%) indicating the efficacy of the sorting/printing process is gentle and has a minimum capacity to introduce systematic biases (see Fig 2G).

**Fig 2.**
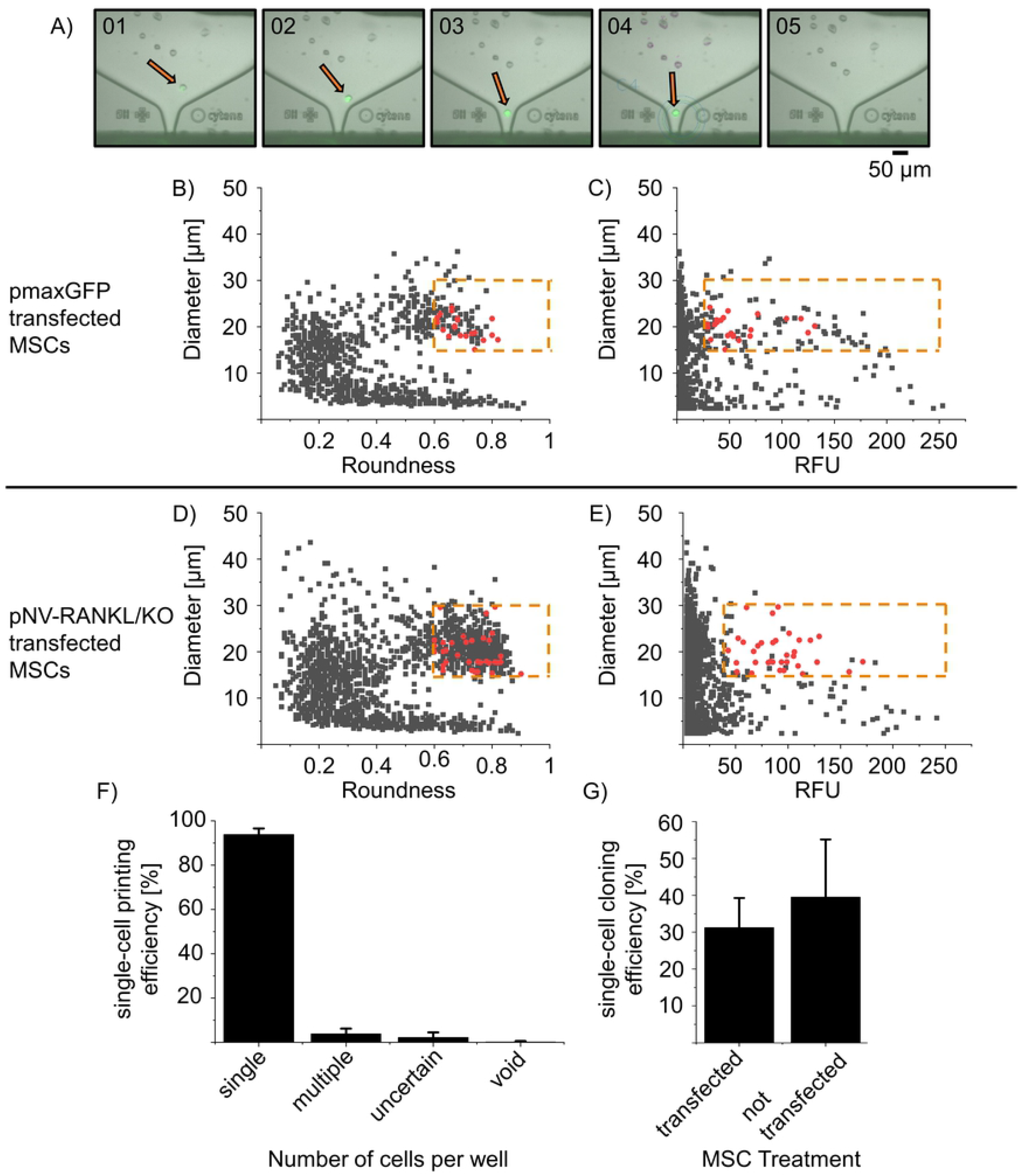
Comparison of the single-cell printing and cloning efficiency of pmaxGFP and pNV-RANKL/KO transfected hTERT MSCs. hTERT MSCs were transfected with either pmaxGFP (transfection efficiency 61%) or pNV-RANKL/KO (transfection efficiency 6%). Two days post-transfection, GFP expressing single-cells were isolated by SCP (see details below). In (A) the SCP (f.sight™) cartridge ejection area is recorded during the printing process, showing a fluorescent single-cell before (01-03), at (04), and after (05) ejection. The SCP algorithm intercepts and deflects cells that are not qualifying the preset parameters (size threshold was set from 15 – 30 *μ*m, roundness from 0.6 – 1, where 1 reflects the perfectly circular object). In (B) and (D) the roundness is plotted against cell diameter and in (C) and (E) the relative fluorescence intensity unit (RFU) is plotted against cell diameter - all detected events are in gray. The red dots represent printed single-cells qualified to the preset parameters (see below). The dotted orange line represents the thresholds of parameters for selecting single cells. (F) The printed single-cells were evaluated for clonality (n= 451 transfected printed cells). The recorded image series of each qualified printing process was evaluated manually, categorizing single, multiple-cell, uncertain, or void printing events. (G) After 2 weeks colonies obtained from single cells were counted and thereby the cloning efficiency determined (n=227 transfected printed cells, n=650 non-transfected printed cells). The single-cell printing parameters applied for pmaxGFP transfected hTERT MSCs: laser intensity of 20 - 40%, an exposure time of 15 - 30 ms and a fluorescence threshold of 30 - 250 RFU while for pNV-RANKL/KO transfected hTERT MSCs laser intensity of 90-95%, exposure time of 80 - 120 ms and a fluorescence threshold of 50 - 250 RFU.

In Fig 2B to E), all recorded events (e.g. debris or living and dead cells) during SCP were plotted against their parameters diameter, roundness, or RFU from one consecutive experiment of either pmaxGFP or pNV-RANKL/KO transfected hTERT MSCs. The cells were sorted in a size range of 15 to 35 *μ*m and roundness of 0.6 to 1, where 1 reflects a perfectly circular object. In Fig 2B and C, the diameter against roundness is plotted while in Fig 2C and D the RFU is plotted against the diameter. The majority of the recorded events show RFU values below 30 but strong population showed an RFU above 30, representing the GFP expressing MSCs with a potential RANKL knockout. The dots marked in red indicate sorted events. The printed cells were selected from an RFU above 30 for pmaxGFP transfected hTERT-MSCs while pNV-RANKL/KO cells were selected from an RFU above 50. In addition, the series of images recorded during the printing process suggest that only cells have been sorted (see exemplarily Fig. 2 A). The selection threshold values are indicated in the graphs by orange dashed lines.

The single-cells were individually sorted and printed in 96-well plates, filled with pre-warmed cell culture medium, and evaluated manually. The first line of analysis is based on the series of images that were taken during the printing process and analyzed manually, whether fluorescent single-cells were successfully sorted and printed or whether multiple, an uncertain amount, no cells or non-fluorescent cells were delivered into the wells. This analysis gives the fluorescent single-cell printing efficiency which is shown in Fig 2F. The combined fluorescent single-cell printing efficiency of all experiments (n=451 transfected printed cells) is 93.9 ± 2.7%, with only 6% of the wells likely containing multiple cells, empty or with uncertain content according to the recorded image series.

These results confirm that the single-cell printer can differentiate morphologically cell-like events from other events and that it can sort cells with different fluorescence intensities since both pmaxGFP transfected hTERT-MSCs with a bright GFP signal and pNV-RANKL/KO transfected hTERT - MSC with a weaker GFP signal were sorted successfully.

The 96-well plates with the isolated single-cells were incubated for 20-25 days and the colonial growth was closely monitored with the NyOne Imager (Synentec). Immediately after printing, the printed cells were detectable in the imager’s FASCC mode due to their GFP signal. After 5 days, the imaging of the cells was switched to bright-field (SCC mode), as the fluorescent signals were lost as expected for transient transfections. Steady colony growth was observed over 20 days without changing the medium (see Fig 3). After 20 days, the numbers of colonies were recorded and the cloning efficiency was calculated relative to the number of printed fluorescent single cells (see Fig 2G). After 20 days, the single-cell colonies were transferred into a cell culture flask and over the course of 120 days, the CPD was measured (for details see materials and methods). The growth rate was recorded for two unmodified hTERT MSC clones (hTERT MSC 1: 0.37 ±0.03 PD/day and hTERT MSC 2: 0.50 ±0.03 PD/day) and one RANKL knockout hTERT MSC clone (g2d: 0.52 ± 0.02 PD/day). All three cell lines show similar growth rates (see Fig 3) so it can be concluded that no significant damage was introduced to the cells by the applied workflow, either through RANKL knockout induced by CRISPR/Cas9 or single-cell isolation with the SCP technology.

**Fig 3.**
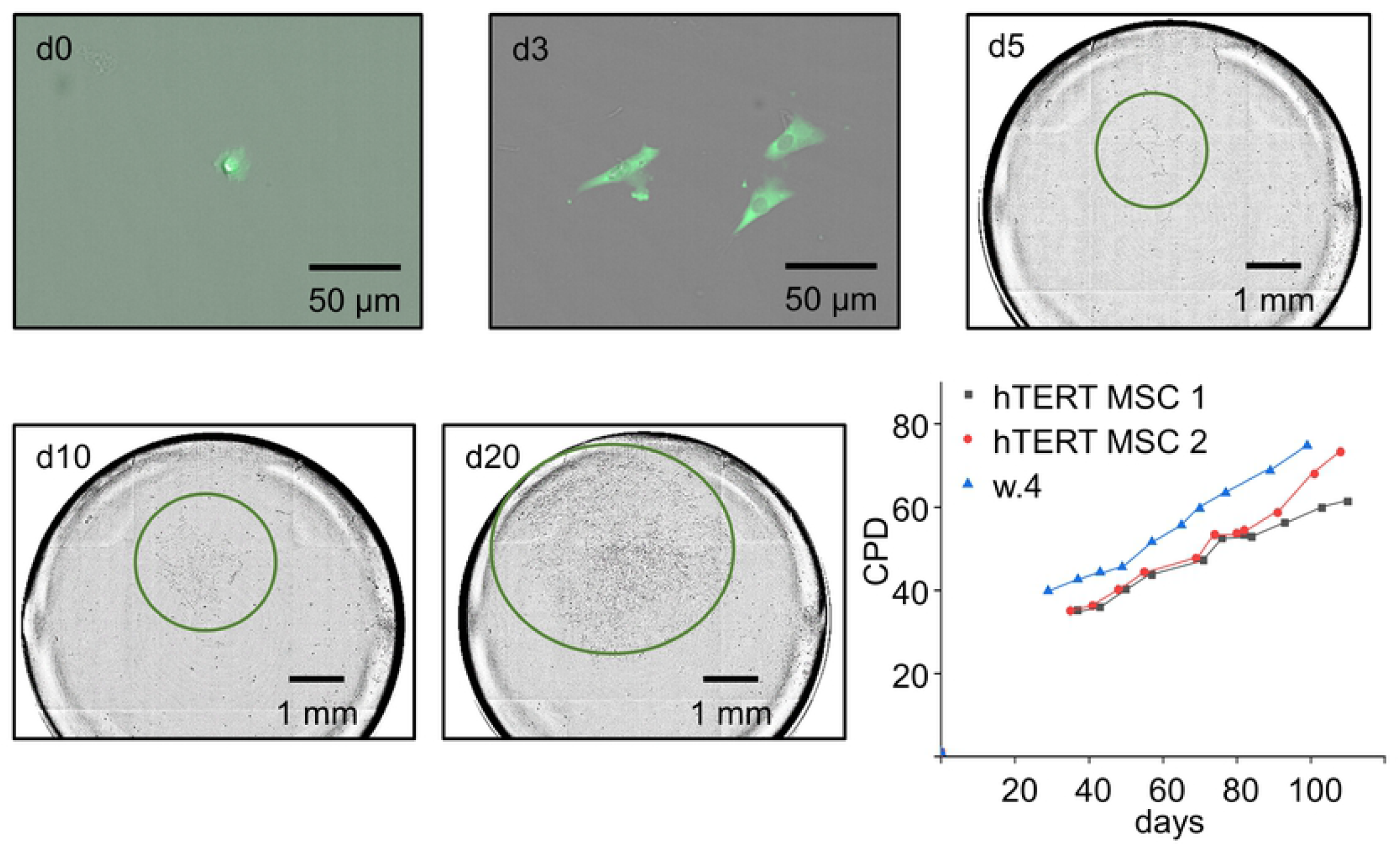
Colony growth of isolated single-cells. The colony growth was recorded with the NyOne™ imager. At day 0 (d0) after printing, the clonality of cells was screened by fluorescent-activated single-cell colony mode (FASCC) and positive wells were marked. During a course of 20 days, FASCC was used until day 3, but from day 5, due to loss of fluorescence, only brightfield single-cell colony mode (SCC) was used and recorded at 5, 10 and 20 days. Colonies are highlighted by green circles. The growth rates of two control hTERT MSC clones and one hTERT MSC RANKL/KO clone (g2d) were tracked for 100 – 120 days after initial single-cell isolation and CPDs are recorded.

### Characterization of the clones

The 96-well plates with the printed, potentially *TNFSF11* edited hTERT MSC clones were cultivated over 20 - 25 days. For subsequent characterization, clones were selected according to their colony morphology and apparent proliferation. 17 were selected for detecting the desired genetic editing. Briefly, all selected hTERT MSC clones were screened for a *TNFSF11* knockout at the genomic level and the edited indel mutations were detected by the surveyor assay (see Fig 4). Control A/B shows three bands at 690 bp, 463 bp and 256 bp for digested heteroduplexes, control A for a homoduplex at 690 bp, genomic DNA from unmodified hTERT MSCs (hTERT) at 1850 bp and the negative control (-) no signal showing the functionality of the assay.

**Fig 4.**
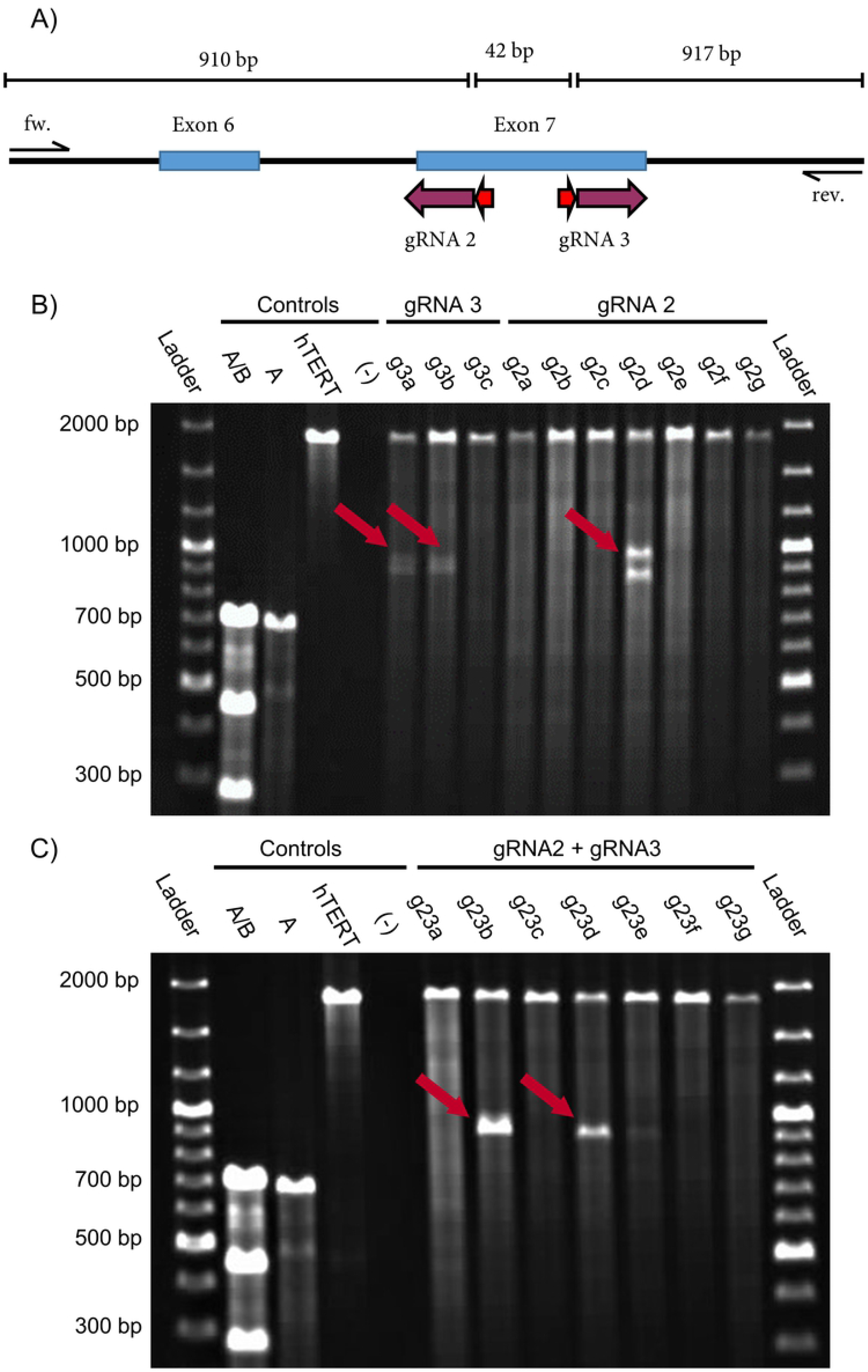
Screening for Indel mutations in genetically modified hTERT MSCs by the surveyor assay. (A) Exons 6 and 7 (blue boxes) from *TNFSF11* with target sites of the corresponding gRNAs (purple arrow). PAM sequences are indicated by red arrows. (B) Agarose gel image (1%) lines are of pNV-RANKL/KO-gRNA2 and pNV-RANKL/KO-gRNA3 transfected hTERT MSCs clones. The clones g3a, g3b and g2d show indel mutations indicated by the red arrows. (C) Agarose gel image of pNV-RANKL/KO-gRNA2 and pNV-RANKL/KO-gRNA3 co-transfected hTERT MSCs clones where clones g23b and g23d show a deletion (red arrows). Controls used: mixed A/B (positive control), pure A (negative control - controls A and B from the kit), unmodified hTERT MSC DNA (hTERT), a template-free negative control (-).

Three bands are expected for the successfully generated clones with different patterns for each gRNA (see Fig 4A). Bands of 1869 bp, 952 bp and 917 bp are predicted for pNV-RANKL/KO-gRNA3, and of 1893 bp, 910 bp and 959 bp for pNV-RANKL/KO-gRNA2. The results of the surveyor assay (Fig 4B) indicate a successful introduction of indel mutation in two out of three pNV-RANKL/KO-gRNA3 transfected hTERT MSC clones (g3a and g3b). On the other hand only one of seven pNV-RANKL/KO-gRNA2 transfected hTERT MSC clones (g2d) showed an alteration at the genetic target site. Cotransfection of gRNA2 and gRNA3 was also attempted in order to delete the genomic DNA between their genomic target sites and therefore hTERT MSCs were co-transfected with the vectors pNV-RANKL/KO-gRNA2 and pNV-RANKL/KO-gRNA3 (see Fig4C). As a consequence of the intended editing, a 42 bp deletion is introduced resulting in the detection by the surveyor assay of the uncleaved band at 1875 bp and two cleaved bands at 910 bp and at 917 bp, respectively. Because these two fragments are of similar size, the pattern for a successful base pair deletion resembles two bands at 1870 bp and 910 bp [38]. The deletion was detected in two of seven hTERT MSC clones (g23b and g23d). Finally, clones g2d, g23b and g23d were selected for further analysis.

As a further step in validating the hTERT MSC *TNFSF11* knockout clones, the gene expression alterations were also investigated. Two exon-spanning qPCRs were used at the exon 4/5- and at the exon 6/7/8-boundaries, respectively. The latter one is located near the site of editing and was used to detect sequences that might be edited, contrary to that the exon 4/5 spanning reaction which was intended to be used as control, as its target site is upstream of the editing site (compare Fig 5C)). *TNFSF11* expression was detected in hTERT MSCs at the CRISPR target site and the upstream exon 4-5 (see Fig 5A and B) [39]. To validate the previous measurements at the protein level, an emulsion coupling reaction was used to confirm the hTERT MSCs RANKL knockout. emulsion coupling uses two oligonucleotide labelled antibodies per target in a homogenous single-molecule assay and determines the absolute counts of the detected ternary complexes. It has a very high femtomolar sensitivity, provides quantitative measurements and is a potential application for high-throughput, automated measurements [20]. Especially, the high sensitivity was crucial as commercial ELISA measurements resulted in obscure data, pointing a necessity in high sensitivity which was further supported by our previous qPCR measurements (LOD ¿10 pg/ml for RANKL see Human TNFSF11 ELISA Kit from abcam #ab213841).

**Fig 5.**
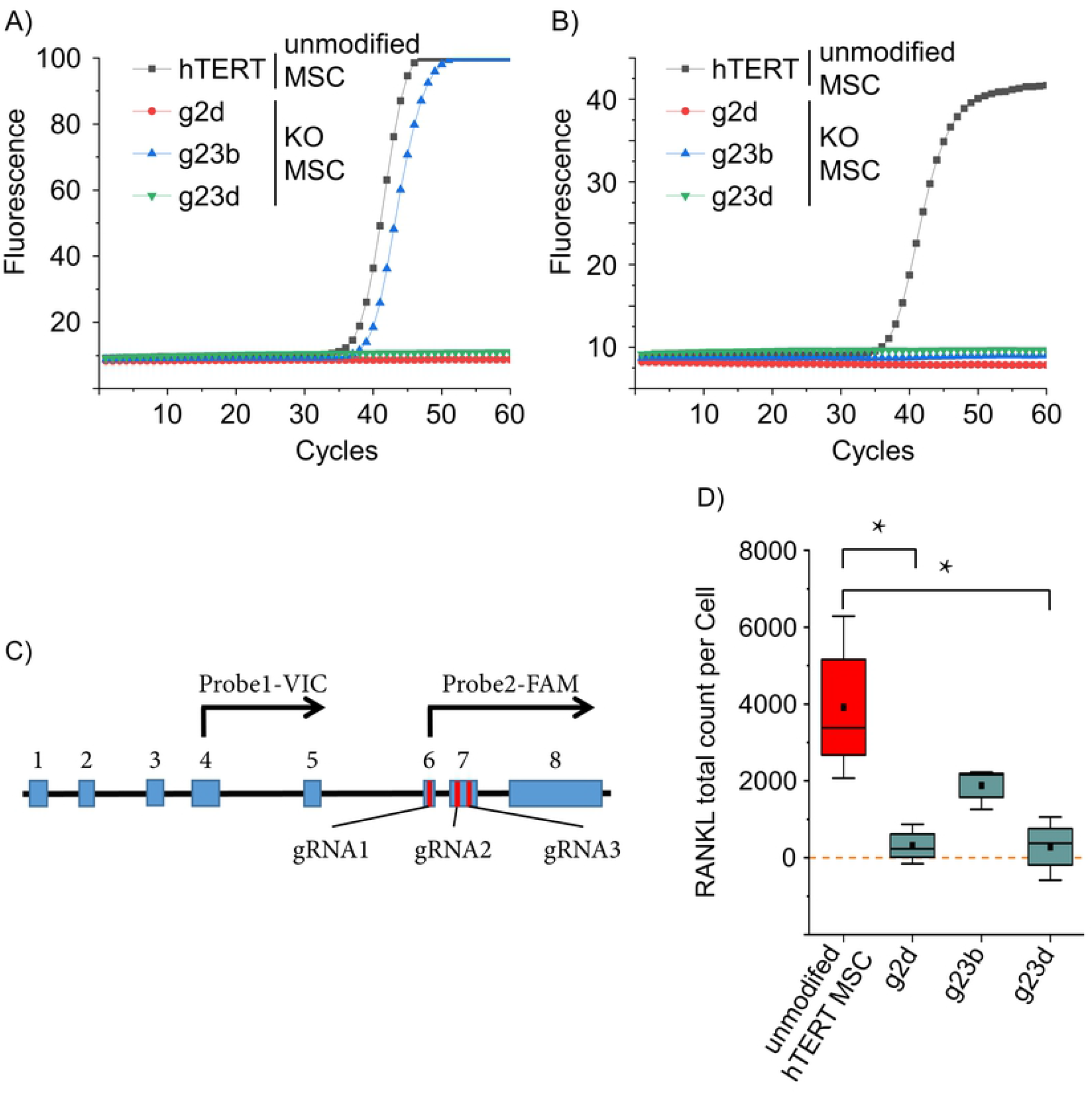
Quantitative measurement of *TNFSF11* RNA and RANKL protein. *TNFSF11* transcription was assessed by two qPCRs upstream (A) and downstream (B) of the gRNAs target sites. hTERT MSC and the *TNFSF11* knockout clones g2d, g23b and g23d were analyzed (n=3, representative runs are displayed). Upstream reaction was positive for hTERT MSCs and g23b, while downstream reaction was positive only for hTERT MSCs. (C) Schematic presentation of the major splice variant of *TNFSF11*. Blue boxes represent exons, red bars indicate the target site of the CRISPR/Cas9 gRNAs and black arrows symbolize the TaqMan probes with the respective fluorophore. (D) Protein expression was measured using emulsion coupling. RANKL-positive hTERT MSCs applied as positive control and 3 different RANKL KO hTERT MSCs were analyzed. The results are in the absolute counts of detected RANKL proteins. The values were normalized against control reaction without interaction (ABC) (red = positive control, cyan = samples); boxes represent the interquartile range (IQR; first to third quartiles); whiskers are 1.5× IQR; horizontal mid-line, median; dot, mean. Statistical significance is indicated: Kolmogorov-Smirnov-test (n = 3) *P ¡ 0.05; zero level means no RANKL protein detected.

The emulsion coupling results in Fig 5D shows the RANKL total count per cell. Originating, non-edited hTERT MSCs were used as a positive control (2897.9 ± 1636.5). RANKL protein expression of clones g2d, g23b and g23d were investigated. Clones g2d and g23d have protein counts of 316.7 (± 520) and 284.6 (± 823.87), respectively. They are not statistically different from zero (One-sample Wilcoxon test, clone g2d P=0.5, n=3; clone g23d P=0.75, n=3) as expected, since genetic alteration was confirmed by surveyor assay and no *TNFSF11* was detected by qPCR pointing towards a complete translational disruption of RANKL. However, hTERT MSC clone g23b has statistically significant RANKL counts of 1882.9 (± 540.22) (see Fig 5D), which is approximately half of the RANKL counts of the positive control (parenteral hTERT MSCs). Interestingly, clone g23b, as well as clone g23d, has a confirmed deletion between gRNA2 and gRNA3, but unlike to clone g23d, the clone g23b has partial positive qPCR results (upstream qPCR positive only) as well, albeit the expression lower (again approximately by a factor of 2) compared to the parenteral hTERT MSCs. We reasonably assume a partial hemiallelic editing of the *TNFSF11* gene and one of the alleles partially remained intact being further supported by both the qPCR and emulsion coupling results (see Fig 5). DNA sequencing could clarify the background of the observed RANKL expression in hTERT MSC clone g23b.

Finally, to investigate the altered differentiation capacity of the edited hTERT MSCs, their intrinsic osteogenic differentiation capacities were tested [35]. The hTERT MSC clone g23d was chosen as it was consistently validated exhibiting biallelic *TNFSF11* knock-out mutations (see above). Its potentially altered differentiation capacity was compared to originated, non-edited hTERT MSCs and confirmed by alizarin red staining after incubation in osteogenic differentiation medium. Cells were incubated over a period of 21 days, with alizarin red staining at day of 7, 14 and 21, respectively. After 14 days, the cells lost their elongated fibroblast-like morphology and began to develop an osteoblast-like morphology and calcified deposits were developed after 21 days, parallel to the parental hTERT MSC control, proving, parallel that the osteoblast differentiation capacity of the edited hTERT MSC clones was not altered during the editing and screening process (see Fig 6).

**Fig 6.**
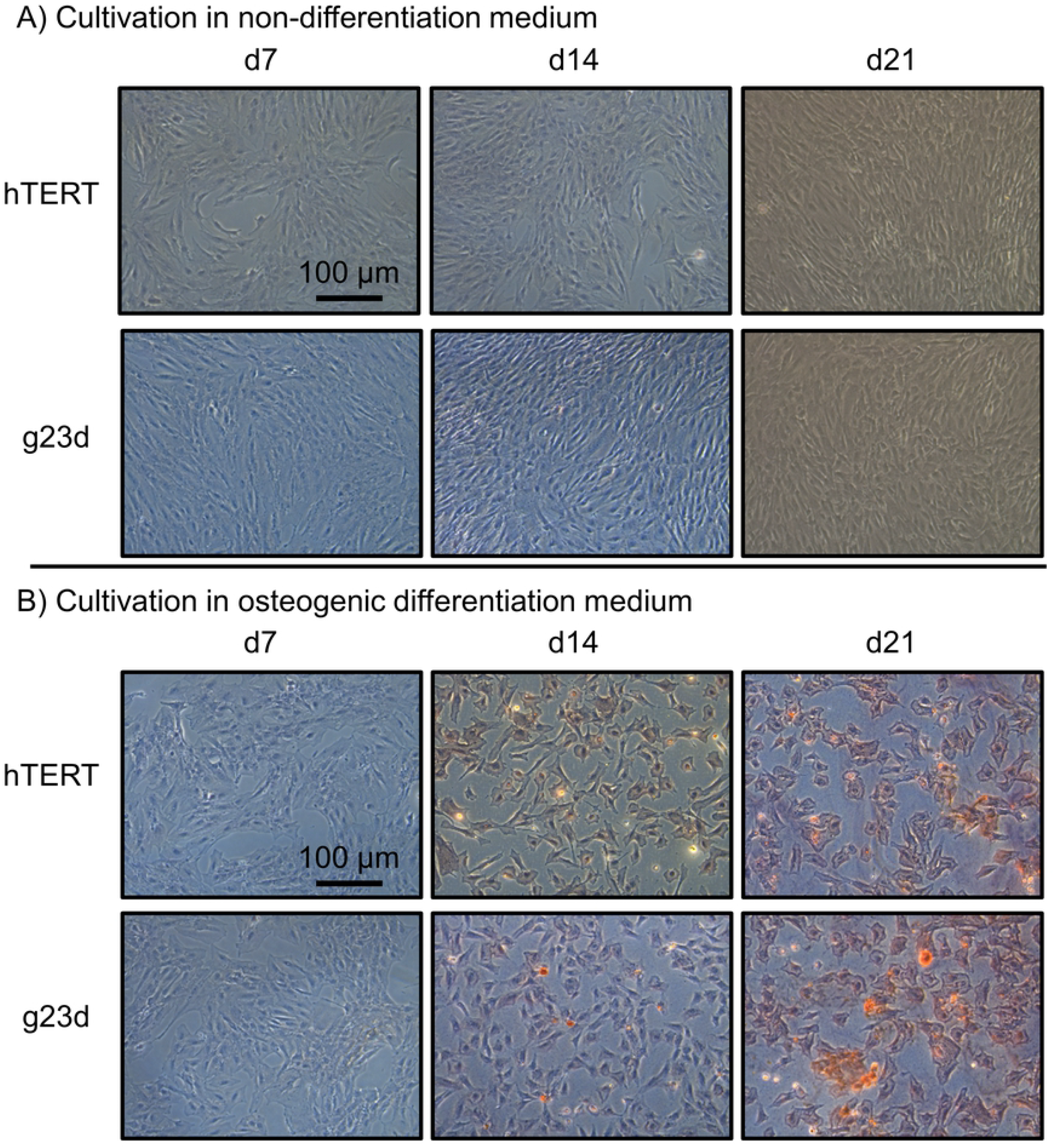
Osteogenic differentiation of hTERT MSCs. Clone g23d along with hTERT MSCs control cells were cultivated in MEM alpha Medium (non-differentiating) and in osteogenic differentiation medium (ODM) in order to induce osteoblast differentiation. Cells in ODM showed osteogenic differentiation, the red staining of cells indicated calcified deposits. Both, hTERT MSCs and clone g23d, readily differentiated into osteoblasts.

## Conclusion

CRISPR-Cas9 is a powerful cell genetic editing tool, but learning and perfecting this revolutionary technology is still advancing [40]. The traditional selection and verification process, despite the CRISPR-Cas9 system’s great efficacy, is still an indispensable part of today’s cloning workflows. Expression of the Cas9 nuclease and a targeted guide RNA (gRNA) in mammalian cells can introduce a double-strand break (DSB) that the cell can repair via non-homologous end joining (NHEJ), an error-prone process that usually results in an insertion or deletion (indel) mutation at the DSB location [41]. The CRISPR-Cas9 system must be optimized both in the gRNA design and transfection efficiency. However, it is equally important to effectively select the generated mixed populations of edited cells, saving both time and resources, while at the same breath ensuring comprehensive and unbiased validation of the edited clones. Here we describe a method to effectively and gently select single-cells for further analysis.

After the introduction of the gRNA(s), single-cells have to be isolated in order to generate clonal lines that can be verified as valid knockouts. Limited dilution cloning and fluorescence-activated cell sorting (FACS) are common, but none is ideal in terms of efficacy, control of selection and cell viability. Limiting dilution cloning is economical and may cause less cellular stress than FACS sorting, but severely limited to generate highly validated single-cell clones. Furthermore, limiting dilution even is introducing affinity-based selection steps and in general the method is still inferior in control [14]. Contrary to that, FACS facilitates excellent control of purity and has high throughput, albeit viability of the sorted cells is a problem. FACS is useful for exploring cell-specific profiles of more than 10,000 cells in each suspension [42]. Many other techniques are also being developed, including manual isolation, microfluidic based sorting mechanisms, magnetic-activated cell sorting, or panning. However, none of them can meet all practical demands [14]. Recently, single cell printing is emerging [18], which is based on an inject-like technology in which free-flying microdroplets that encapsulate cells are generated by a non-contact dispensing procedure. SCP uses an automated visual recognition of cells, brightfield and/or fluorescent, to confirm the presence of a single cell in the next ejected droplet. It has limited fluorescence sorting capabilities, but is generally suitable for cloning applications. The ejected microdroplet of 150 pL can be deposited on various substrates, and the printing process of each ejected cell is recorded. The method is fast, very gentle on cells, has a throughput comparable to FACS and provides an ¿99.99% assurance that the cell lines derived from this workflow have been clonally derived [43].

The SCP was used to sort GFP-gRNA transfected hTERT MSCs. According to the results, the transfected cells were clearly distinguishable from the non-transfected background cells, having fluorescent single-cell printing efficiency of 93.9% (± 2.7%). It delivered a cloning efficiency of 31.3% (±8%), which was not statistically different from the cloning efficiency of non-transfected hTERT MSCs subjected to the same process. These excellent sorting capabilities are also vital at low transfection efficiencies, and demand less optimization of transfection protocols, eliminating a significant bottleneck in large-cell cloning efforts. The cloning process was also monitored by automated microscopy, estimating the population doubling levels (PDL) of the cell clones. Although it varied slightly for each clone, it also showed no significant difference - around half doubling per day - for both transfected and non-transfected hTERT MSCs. As a conclusion, SCP has a minimal impact on cell viability, as also confirmed in the osteogenic differentiation assay, and is superior over the other techniques in many aspects. The ease of use, including the use in sterile operational environments, compatibility with various downstream processes, and the wealth of verification data, makes it ideal for single-cell cloning application, even in high-demand commercial environments.

The confirmation of valid knock-outs is a tedious process because many levels of verification are mandatory, including genetic screening for homozygous/heterozygous knock-outs, transcription, and protein expression. In large-scale characterization efforts of single-cell clones, the notable bottleneck is the necessary clonal expansion, especially for clonal protein analytics. Since less sample demanding techniques for the genetic (PCR based using deletion exon-spanning primers, surveyor assay or sequencing) and transcriptomic (RT-PCR) analysis are readily available. Protein analytics is lagging behind on those PCR based techniques especially in sensitivity, which renders a protein analytically proof of a knock-out very hard to establish, especially in the case of low abundance proteins. In our experiments the detection of RANKL by ELISA has failed with an assay LOD of ¿ 10 pg/ml (see Human TNFSF11 ELISA Kit from abcam #ab213841). Contrary to that, the molecular sensitivity of emulsion coupling enables low sample amounts down to a few single proteins. Single-cells can be assayed projecting a possible economical large-scale characterization of the single-cell clones at the protein level. However, the introduction of emulsion coupling as a general element of clone characterization workflows offers many advantages apart from low samples sizes. emulsion coupling can be parallelized to detect many proteins in a single assay. Thus, it is envisaged that the functional consequences of the knock-outs can also be assayed, which opens up a new direction in clone verification. Especially, since emulsion coupling can detect the interactions and post-translational modifications of the proteins simultaneously. Regarding our results using emulsion coupling, we could confirm the RT-PCR based hemizygosity of the introduced deletion by detecting a lower than wild-type amount of RANKL in the case of clone g23b. We could also show that the homozygous knock-out clones were clearly negative in this assay. These quantitatively congruent results can also confirm the overall integrity of the emulsion coupling measurements.

In summary, this workflow can be adopted for other cell types and genetic targets and has great potential for many applications in which monoclonal cells are required. There are no restrictions for the single-cell printing technology to handle different cell types [44], and emulsion coupling can reduce both the required time and resources and potentially augment the content of the characterization workflows, enabling better clones with lower failure rates in the late stages of cell-line developments.

## Supporting information

**Fig 7. S1_raw_images.** Raw gel images which were used for Surveyor Assay.

## Acknowledgments

The authors thank Prof. Dr. Matthias Schieker from the Laboratory of Experimental Surgery and Regenerative Medicine of the Ludwig-Maximilians-University of Munich for providing the immortalized hTERT MSC cell line. This work was supported by funding of the Bundesministerium fiir Bildung und Forschung (GA 031B0114C).

